# Master Transcription Factors Regulate the DNA Methylation Landscape During Hepatocyte Differentiation

**DOI:** 10.1101/2020.12.16.423165

**Authors:** Takahiro Suzuki, Erina Furuhata, Shiori Maeda, Mami Kishima, Yurina Miyajima, Yuki Tanaka, Joanne Lim, Hajime Nishimura, Yuri Nakanish, Aiko Shojima, Harukazu Suzuki

## Abstract

**Background:** Hepatocytes are the dominant cell type of the human liver, with functions in metabolism, detoxification, and in producing secreted proteins. During the process of hepatocyte differentiation, gene regulation and master transcription factors have been extensively investigated, whereas little is known about how the epigenome is regulated, particularly the dynamics of DNA methylation, and the upstream factors that have critical roles.

**Results:** By examining changes in the transcriptome and the methylome during *in vitro* hepatocyte differentiation, we identified putative DNA methylation-regulating transcription factors, which are likely involved in DNA demethylation and maintenance of hypo-methylation in a differentiation stage-specific manner. Of these factors, we further reveal that GATA6 induces DNA demethylation together with chromatin activation at a binding-site-specific manner during endoderm differentiation.

**Conclusions:** These results provide an insight into the spatiotemporal regulatory mechanisms exerted on the DNA methylation landscape by transcription factors, and uncover a new role for transcription factors in early liver development.

## Background

Hepatocytes, the major parenchymal cells in the liver, are responsible for key liver functions such as metabolism and detoxification. In embryogenesis, the first fate decision to the hepatocyte lineage is the differentiation of primitive streak cells to definitive endoderm (DE) cells, which are a common precursor of endoderm tissues such as liver, pancreas, and gut. Hepatoblasts are hepatic progenitor cells derived from the DE cells, which then differentiate into fetal-like hepatocytes and mature hepatocytes in a stepwise manner. Thus, hepatocytes emerge from pluripotent stem cells through several progenitor cell types.

Several transcription factors (TFs), including c-Jun, and members of the HNF and GATA families are known to play important roles in liver development and hepatocyte differentiation [1–9]. For instance, transcription factor HNF4A is indispensable for specification and early development of the liver [9]. Furthermore, HNF4A is required for the transcriptional activation of genes such as CYP3A4 and CYP2D6, which are crucial for hepatocyte metabolic functions [3,4]. GATA4 and GATA6 are also essential for development of endoderm-derived tissues and cells, including hepatocytes [6–8]. Notably, GATA6 knock-out mice die around E5.5 due to a deficiency of extra-embryonic endoderm development, which can be rescued by tetraploid embryo complementation assays, and indicating that GATA6 is required for liver development and hepatic specification [8,10–12]. Thus, multiple TFs sequentially and coordinately regulate peripheral genes necessary for hepatocyte differentiation.

Gene expression dynamics are regulated not only by the action of transcription factors but also by epigenetic modifications such as DNA methylation. In mammals, most DNA methylation occurs at the cytosines of CpG dinucleotides, adding a methyl group at the 5-carbon of the cytosine. DNA methylation of gene regulatory regions appears to be associated with silencing of the expression of the downstream gene [13]. Specifically, gene regulatory regions must be demethylated for activation of the downstream gene. Consistent with this, the DNA methylation profile is dramatically altered during embryogenesis and cellular differentiation, with roles in tightly regulating expression of downstream genes [14–17]. Indeed, it is reported that DNA methylation plays a crucial role in the expression of numerous liver-specific genes [12,18–24].

Furthermore, expression of CEBP hepatic transcription factors are affected by treatment with the DNA methyltransferase (DNMT) inhibitor, 5-Aza-dC[25]. Interestingly, the DNMT inhibitor facilitates trans-differentiation of adipose tissue-derived stem cells or mesenchymal stem cells to hepatocyte-like cells[26–28]. Collectively, these findings show that DNA methylation is a crucial factor for hepatic differentiation.

The gain of DNA methylation is directly achieved by *de novo* DNMTs [17,29–31], and methylation status is maintained during cell divisions by a maintenance DNMT [31–34]. If DNA methylation maintenance does not work properly, the level of DNA methylation declines upon cell proliferation, which is known as passive DNA demethylation[35]. Alternatively, it is plausible that sequential oxidative processes achieve active DNA demethylation by ten-eleven translocation (TET) enzymes [36–40], followed by base-excision repair[40,41]. In addition, the oxidized forms of methylated cytosine (5-hydroxymethyl cytosines (5hmC), 5-formyl cytosine (5fC), and 5-carboxy cytosine (5caC)) are also depleted by passive demethylation mechanisms, because these bases are not recognized by the maintenance DNA methylation mechanism [42,43]. Thus, DNA methylation is a balance between gain and loss of methylated bases.

In addition to the mechanisms by which DNA methylation is gained and lost, mechanisms underlying spatiotemporal regulation of DNA methylation are also critical in understanding the overall dynamics of DNA methylation. We and other groups recently reported that some TFs regulate the timing and site-specificity of DNA demethylation [44–50]. We found that RUNX1, an essential transcription factor for hematopoietic development and immune cell functions, induces DNA demethylation by recruiting the TET enzymes and TDG to their binding sites[44]. We have also identified eight novel DNA-demethylating TFs using a screening method we developed[45]. In addition to our findings, other groups have reported DNA-demethylating TFs with roles in several biological processes[46–50]. Thus, a growing body of evidence suggests critical roles for TFs in the regulation of DNA methylation. However, the epigenetic roles of TFs specific for hepatocyte differentiation have yet to be identified.

In the present study, we combine TF binding motif (TFBM) overrepresentation analysis for differentially methylated regions [45] with transcriptome analysis. We identify TFs with putative roles in regulating DNA methylation during hepatocyte differentiation by studying *in vitro* the process of hepatocyte differentiation from human induced pluripotent stem (iPS) cells. Of these TFs, we validate that GATA6 is a master regulator for both DNA demethylation and chromatin activation in the differentiation of the DE. Our data provide significant insights into the regulatory mechanisms shaping the DNA methylation landscape during hepatocyte differentiation.

## Results

### DNA methylation dynamics throughout hepatocyte differentiation

We induced hepatocytes from human iPS cells *in vitro* and examined the transcriptome by Cap Analysis Gene Expression (CAGE)[51] (Fig. 1A, B). Expression of pluripotent marker genes *(POU5F1* and *NANOG*) was considerably downregulated after day 7 of differentiation and undetectable after day 14 (Fig. S1). In contrast, DE markers (*SOX17* and *FOXA2*) and hepatic markers (*HNF1B*, *PPARA*, *AFP*, and *PAX6*) were upregulated at day 7 and day 14-to-day 28, respectively (Fig. S1). Notably, because *AFP* is known to be upregulated in immature hepatocytes and to be downregulated in mature hepatocytes, and *PAX6* is a maturation marker of hepatocytes, our data confirmed the *in vitro* differentiation mimics the whole process of *in vivo* hepatocyte differentiation[52]. Thus, our time-course samples represent day 0 as iPS cells, day 7 as DE, day 14 as hepatoblasts, day 21 as fetal-like hepatocytes, and day 28 as mature hepatocytes, respectively (Fig. 1A, B).

**Fig. 1.**
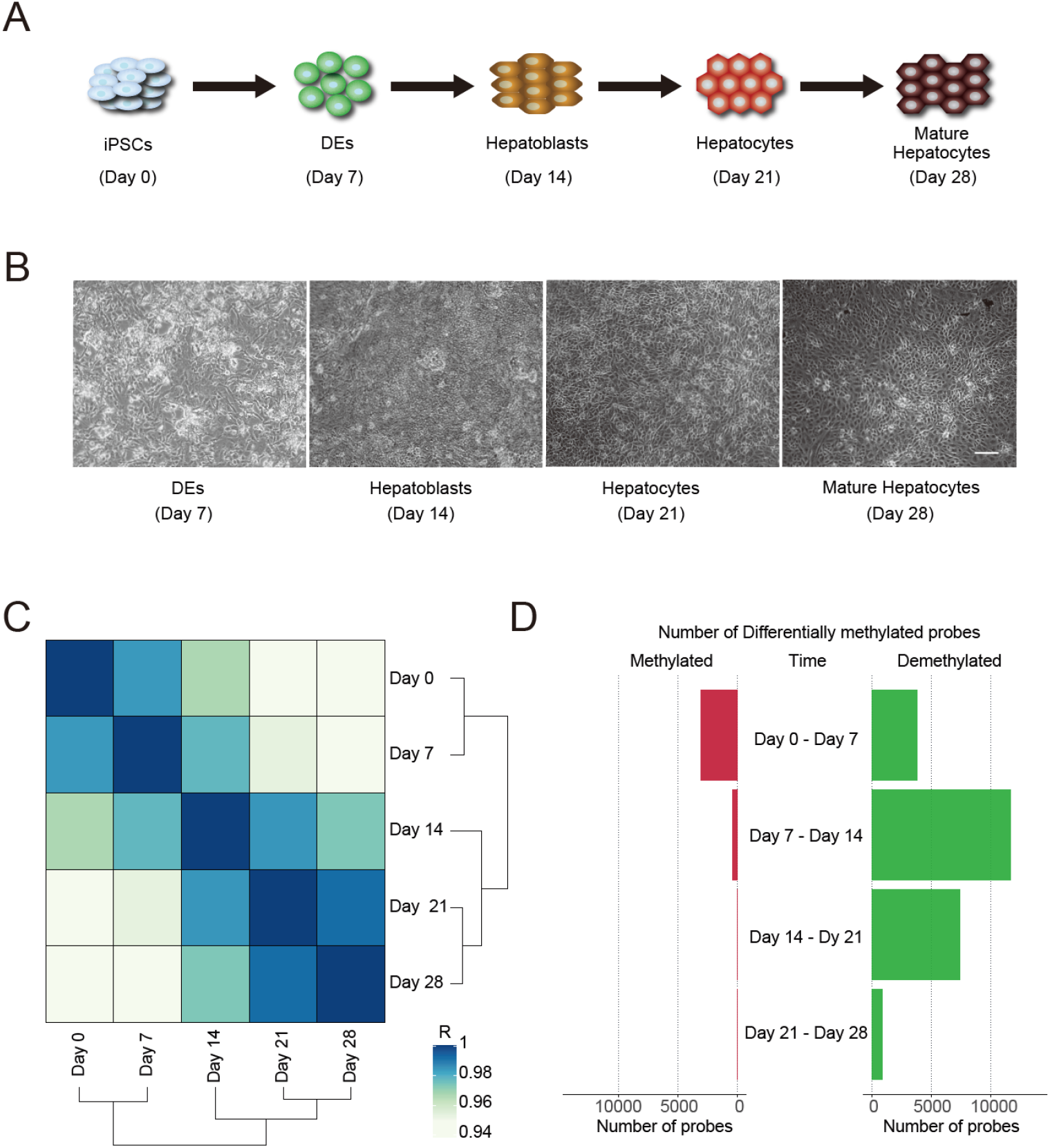
Time-course methylome analysis of hepatocyte differentiation. **(A)** Schematic illustration of *in vitro* hepatocyte differentiation. **(B)** Pictures of each time point. The scale bar is 200 μm. **(C)** A correlation matrix with hierarchical clustering. The color represents the correlation coefficient (R). **(D)** The number of differentially methylated probes. The left bar plot (red) is methylation and the right bar plot (green) is demethylation.

To investigate changes in DNA methylation during hepatocyte differentiation, we performed a methylome analysis of the time-course samples using MethylationEPIC BeadChip (Illumina). Hierarchical clustering showed that iPS cells and DE cells were segregated from the differentiated cells that followed in the time-course, consistent with a commitment to the hepatocyte lineage (Fig. 1C). Comparing adjacent time points, we identified 3088, 446, 38, and 54 methylated CpGs and 3809, 11652, 7383, and 864 demethylated CpGs in each interval (Fig. 1D). Thus, although the gain of methylation mostly occurs in early time points, the number of the differentially methylated CpGs were biased toward demethylation in all intervals, indicating that demethylation (loss of methylation) predominates in the dynamics of DNA methylation during hepatocyte differentiation.

We associated biological functions to the differentially methylated regions using the Genomic Region Enrichment of Annotations Tool (GREAT)[53] and summarized the results based on semantic similarity[54]. This analysis revealed an enrichment in development and morphogenesis related Gene Ontologies (GOs), including “pattern specification process”, “anatomical structure development”, “radial pattern formation”, “developmental process”, and “regulation of developmental process” (Fig S1B and C). Overall, these results imply that DNA methylation mainly regulates genes related to the developmental process, consistent with specifying the cells into the hepatocyte lineage.

### Prediction of DNA methylation-regulating transcription factors throughout hepatocyte differentiation

We previously developed a screening system to identify TFs which regulate binding site-directed DNA methylation (hereinafter referred to as DNA methylation-regulating TFs), which is based on TF binding motif (TFBM) overrepresentation analysis for differentially methylated CpG regions using ectopic TF overexpression [45]. By modifying this system, we here performed TFBM overrepresentation analysis for the differentially methylated CpG regions between two adjacent time points of the differentiation time-course with the TFBM position weight matrix (PWM) database of the IMAGE tool [55]. This database covers most of the known TFs. Because some TFs, such as TFs in the same family, share the same or similar binding motif, the results of TFBM overrepresentation analysis often include false positives. Therefore, to reduce the possibility of false positives, we further narrowed down the overrepresented TFBMs by considering TF expression (CAGE tag per million (TPM) ≥ 50) in either of the two adjacent time points of an interval (Fig. 2A). Thus, by combining methylome and transcriptome analyses, we identified putative DNA methylation-regulating TFs. Comparing each adjacent timepoint, we identified in total 16 putative DNA methylation-regulating TFs in the methylated regions. Of these, 13 TFs, including POU5F1, a pluripotent cell-specific TF, were identified in the DE differentiation stage (Day 0 -to- Day 7) (Fig. 2B). In addition, GATA6, GATA3, and GATA4 were identified in the hepatoblast differentiation stage (Day 7 -to- Day 14) (Fig. 2B). Interestingly, these putative DNA methylation-regulating TFs for the methylated regions were prone to being highly expressed in the earlier time point of the intervals and then declined along with the progress of differentiation (Fig. 2C and D; Fig. S2A).

**Fig. 2.**
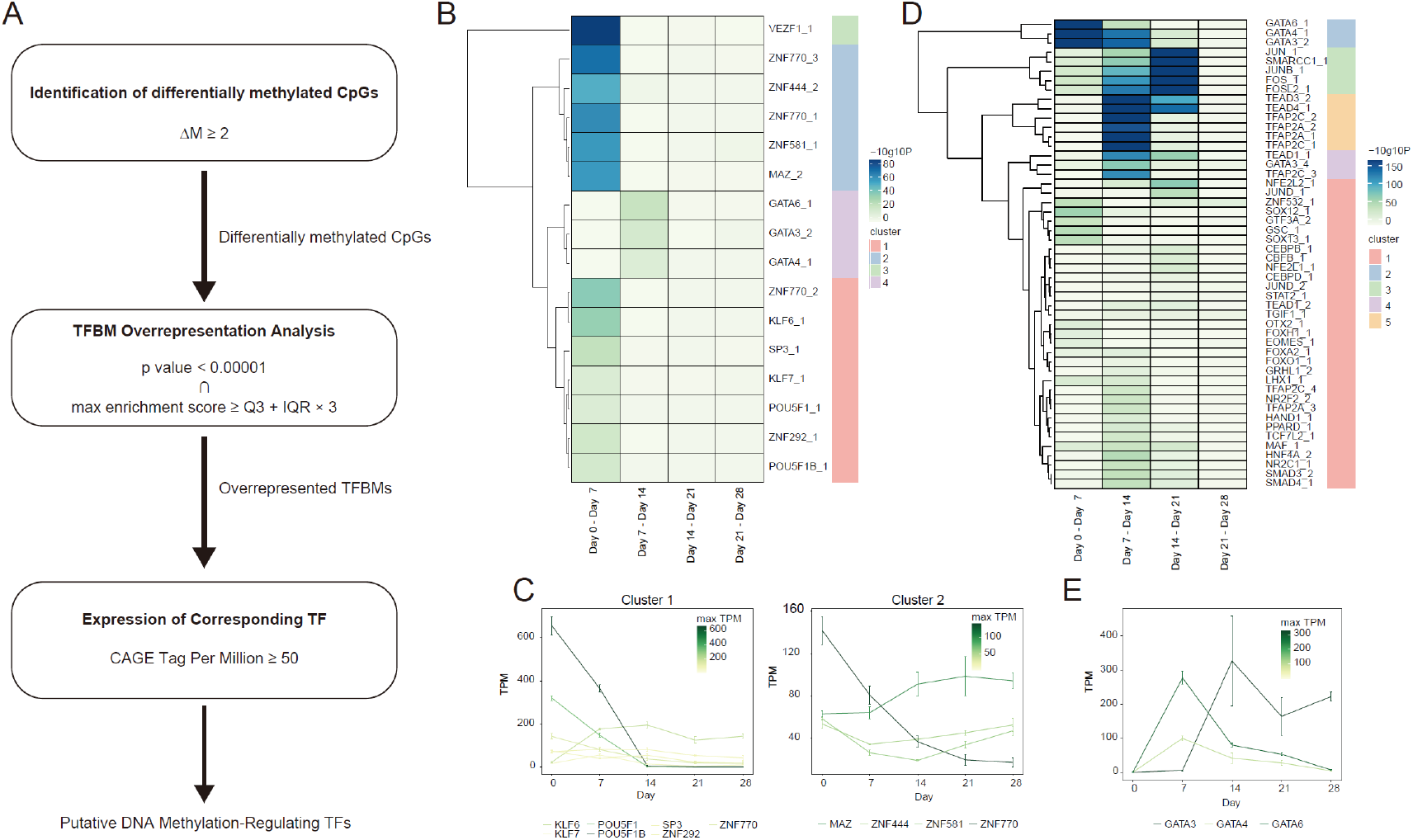
Prediction of DNA methylation-regulating TFs. **(A)** The workflow of DNA methylation-regulating TF prediction. (B, D) Heatmap showing the *p*-value of over-represented TF binding motifs at methylated **(B)** and demethylated **(D)** regions. Each column is an interval of adjacent time points. Each row is a putative methylation-regulating TF. Dendrogram of hierarchical clustering is shown at the left of the heatmap and clusters are shown at the right of the heatmap as colors. **(C)** mRNA expression profile of the cluster 1 and 2 putative DNA methylation-regulating TFs for methylated regions. X- and Y-axes show time points of differentiation (hours from differentiation initiation) and tag-per-million (TPM) of CAGE, respectively. The color of each line represents the maximum TPM. Cluster 3 and 4 and the putative DNA methylation-regulating TFs for demethylated regions were shown in Fig. S2 **(E)** mRNA expression profile of the GATA3, GATA4, and GATA6. X- and Y-axes show time points of differentiation (hours from differentiation initiation) and tag-per-million (TPM) of CAGE, respectively. The color of each line represents the maximum TPM.

On the other hand, we identified 50 putative DNA methylation-regulating TFs at demethylated regions. Of these, HNF4A, an essential TF for liver development [3][4], was identified in the hepatoblast differentiation stage (Fig. 2D). In addition, the overrepresentation of TFBMs for Activator Protein 1 (AP-1) components such as JUN and FOS, which are involved in the stress response and regeneration in the liver[56–58], increased from the DE differentiation stage to the fetal-like hepatocyte differentiation stage (day 14 -to-day 21) (Fig. 2D). Importantly, GATA6, GATA4, and GATA3, which were also identified in the methylated regions of the hepatoblast differentiation stage, were firstly identified in the DE differentiation stage and overrepresentation of these binding motifs declined as differentiation proceeded (Fig. 2D). Contrary to the putative DNA methylation-regulating TFs for the methylated regions, expression of the putative DNA methylation-regulating TFs for the demethylated regions tends to be upregulated in later timepoints of the intervals (Fig. 2E; Fig. S2B). Taken together, these results suggest that diverse TFs cooperatively regulate the DNA methylation landscape. In particular, GATA transcription factors appear to be the major factors for the DNA methylation regulation, participating in both methylation and demethylation changes.

### Ectopic GATA6 overexpression induces binding site-directed DNA demethylation

Our data suggested the GATA family is a crucial factor for regulating DNA methylation during hepatocyte differentiation, mainly contributing to the demethylation that occurs in DE differentiation. Of the GATA proteins, GATA4 and GATA6 were consistent with the pattern of mRNA expression and are known to be essential TFs for the DE differentiation stage [6–8]. Therefore, we focused the following analysis on possible epigenetic functions of GATA4 and GATA6 in DE differentiation. Firstly, we performed qRT-PCR to confirm the expression changes of *GATA4* and *GATA6* during the DE differentiation stage. Expression of both *GATA4* and *GATA6* started increasing from 48 hours after induction of the differentiation and were maximized at 66 hours and 60 hours, respectively, suggesting that *GATA6* expression precedes *GATA4* expression (Fig. 3A).

**Fig. 3.**
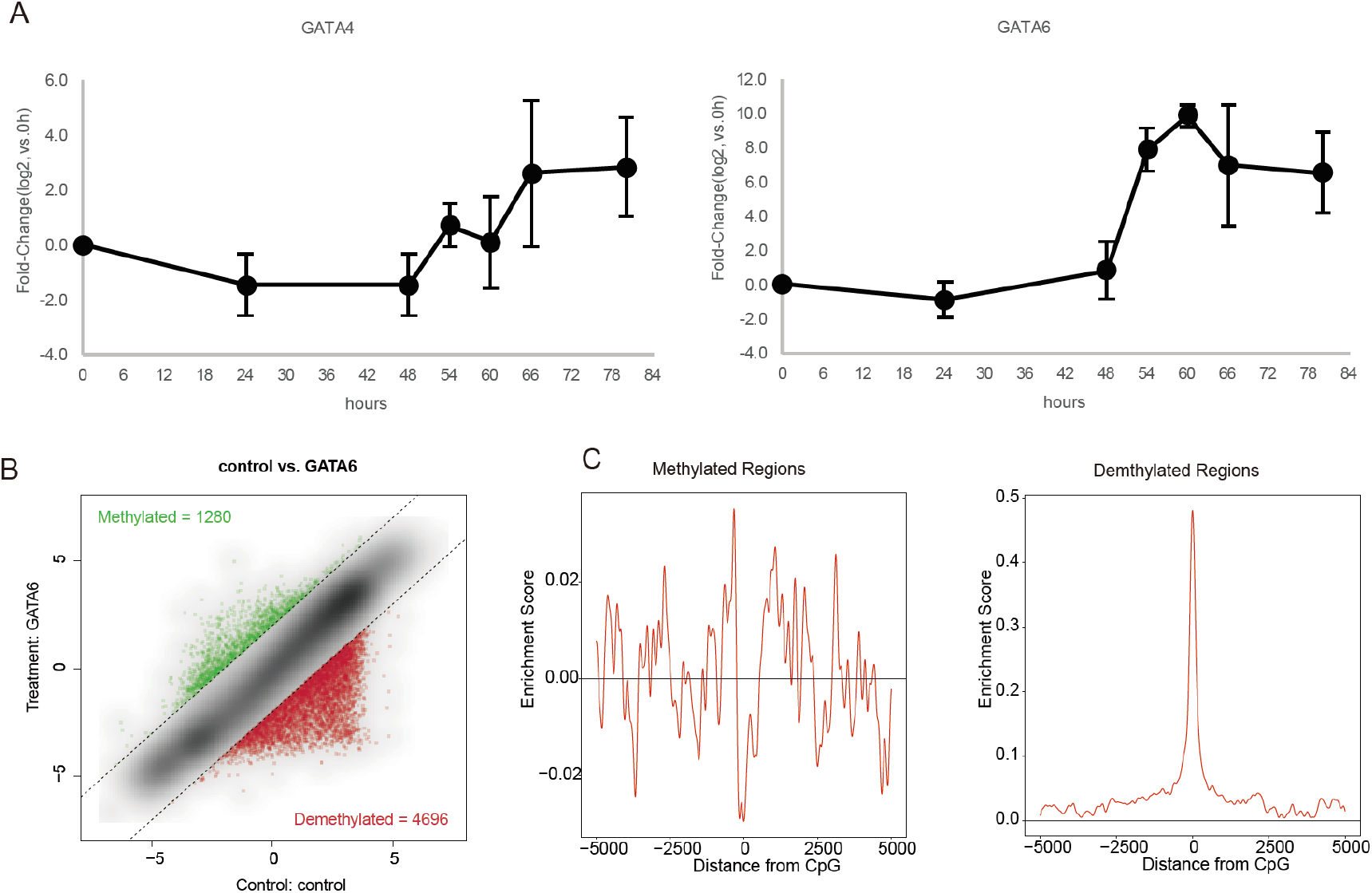
GATA6-mediated binding site-directed DNA demethylation. **(A)** qRT-PCR analysis for GATA4 (left) and GATA6 (right). X- and Y-axes show time points of differentiation (hours from differentiation initiation) and fold-change (compared with 0 hours, log2 scaled), respectively. (**B**) Scatter plot showing M-value of each probe. X- and Y-axes show M-values of the control sample and GATA6-overexpressing sample, respectively. Dotted lines represent ΔM = 2 or −2. Green and red dots are methylated and demethylated probes, respectively, and the number of each probe is shown at the upper left and lower right. (**C**) Distribution of enrichment score for the GATA6 binding motif within ±5,000 bp of methylated (left) and demethylated (right) CpG probes in GATA6-overexpressing 293T cells. X- and Y-axes show distance from probe CpG position and enrichment score, respectively. Horizontal lines are enrichment score = 0.

Furthermore, *GATA6* expression increased drastically, greater than 1,000-fold at 60 hours compared with 48 hours, whereas *GATA4* expression increased only 4-fold at 66 hours compared with 48 hours, indicating the dominant impact of GATA6 (Fig. 3A). Indeed, GATA6 is reported to be an upstream factor of GATA4[11]. Therefore, we next overexpressed GATA6 in HEK293T cells, followed by methylome analysis to investigate the role of GATA6 in regulating DNA demethylation. In the HEK293T cells overexpressing GATA6, we identified 1,280 and 4,696 methylated and demethylated CpGs, as compared with mock control transduced cells (Fig. 3B). The motif overrepresentation analysis for the differentially methylated regions revealed that the GATA6 binding motif was significantly overrepresented at the demethylated regions but not at methylated regions in the GATA6 overexpressing cells, indicating that GATA6 functions in binding site-directed DNA demethylation (Fig. 3C).

### DNA demethylation accompanies GATA6 binding during iPS-DE differentiation

To investigate the dynamics by which GATA6 regulates DNA demethylation, we performed finer time-course transcriptome and methylome analyses during the time-window of GATA6 emergence (after 0 hours (h), 48 h, 54 h, 60 h, 66 h, and 72 h of the differentiation process) (Fig. 4A). T, a marker of the primitive-streak, was upregulated at 48 h and was downregulated after 54 h. DE markers were upregulated during the period 48 h-to-72 h (Fig. S3A). In agreement with the qRT-PCR analysis (Fig. 3A), the expression of GATA6 was slightly upregulated at 48 h and drastically increased after 48 h (Fig. S3A). Hence, our data indicate DE commitment occurs during the period 48 h - to- 72 h into the differentiation process.

**Fig. 4.**
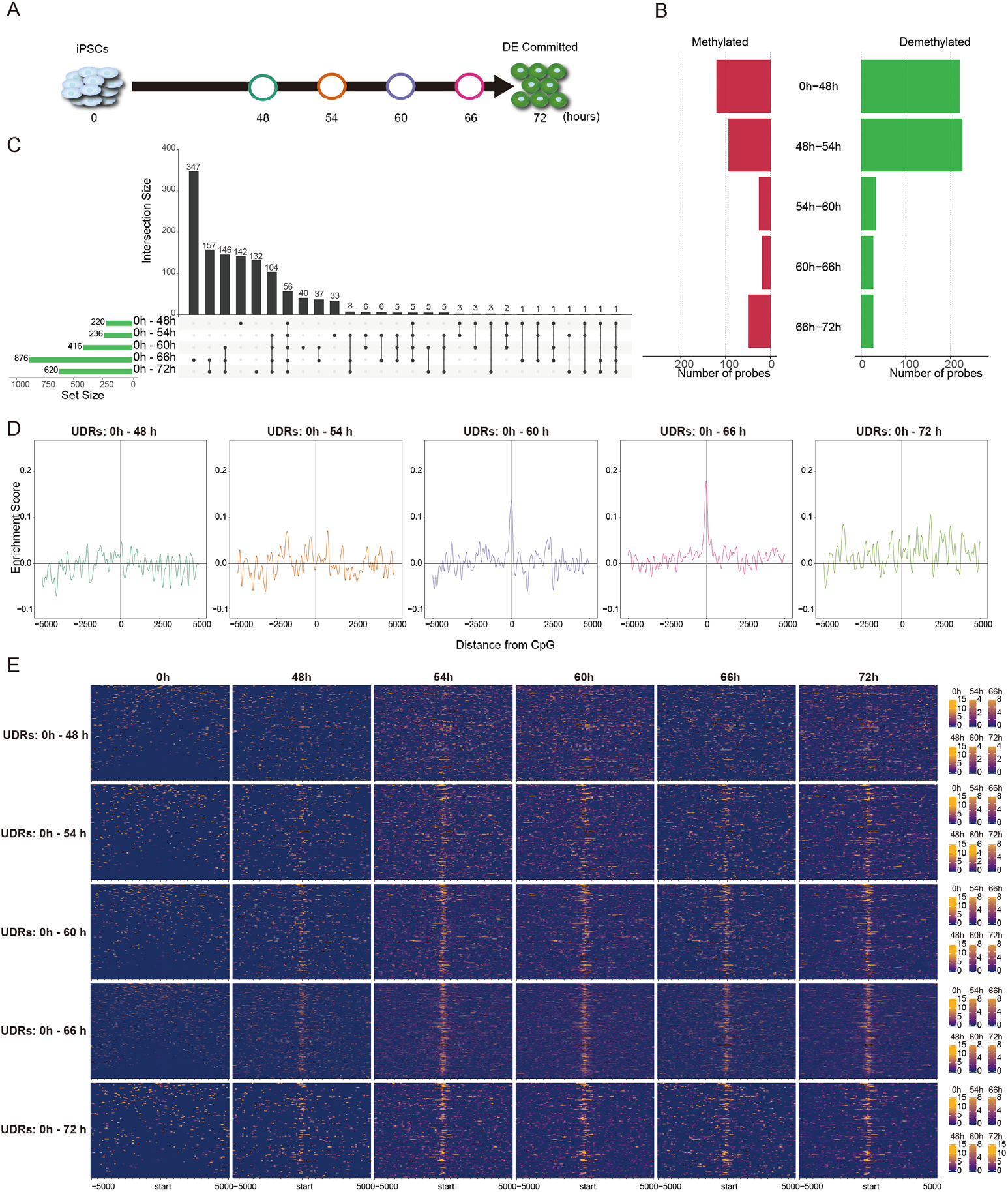
GATA6-mediated DNA demethylation analysis during DE differentiation. **(A)** Schematic illustration of time-course sampling of DE differentiation. **(B)** The number of differentially methylated probes. The left bar plot (red) is methylation and the right bar plot (green) is demethylation. **(C)** UpSet plot showing the demethylated probes at each comparison. The vertical bars indicate the number of intersecting demethylated probes between comparisons, denoted by the connected black circles below the histogram. The horizontal bars show the demethylated probe set size. **(D)** Distribution of enrichment score for the GATA6 binding motif within ±5,000 bp of demethylated CpG probes at each time point compared with undifferentiated iPS cells (0 hours). X- and Y-axes show distance from probe CpG position and enrichment score, respectively. Horizontal and vertical lines are enrichment score = 0 and demethylated CpG position, respectively. The colors of each plot represent colors of timepoints shown in Fig. 4A. **(E)** Enrichment heatmap showing coverage of GATA6 ChIPmentation reads at a range of ± 5 kbp from demethylated CpGs. Each time point is horizontally aligned and each of the UDRs are vertically aligned. Dark blue is low coverage and orange is high coverages.

By comparing adjacent time points, we identified 120 (0 h -to- 48 h), 94 (48 h - to- 54 h), 26 (54 h -to- 60 h), 19 (60 h -to- 66 h), and 50 (66 h -to- 72 h) methylated CpGs and 220 (0 h -to- 48 h), 226 (48 h -to- 54 h), 33 (54 h -to- 60 h), 27 (60 h -to- 66 h), and 27 (66 h -to- 72 h) demethylated CpGs, respectively (Fig. 4B). However, we did not find the GATA6 binding motif overrepresented at those demethylated regions during any interval (Fig. S3B). Because the time intervals between adjacent time points are 6 hours except for the initial period (0 h -to- 48 h), the changes in methylation levels may not be enough to be detected as demethylation (ΔM > 2). Indeed, the GATA6 binding motif was overrepresented at the regions demethylated between 0 h and 72 h and these demethylated regions tend to be continuously demethylated from 0 h (Fig. S3C and D). Therefore, to investigate whether the GATA6 binding motif is overrepresented for the cumulative changes in methylation, we compared the regions demethylated at each time point with that at 0 h. We identified 220 (0 h -to- 48 h), 236 (0 h -to- 54 h), 416 (0 h -to- 60 h), 876 (0 h −66 h), and 620 (0 h -to- 72 h) demethylated-CpGs (Fig. 4C). Because these demethylated CpGs include those that were demethylated in the earlier time point that were newly detected as demethylated CpGs at each timepoint (referred to as uninherited demethylated CpGs) to clarify the effects of each additional time period. GATA6 motif overrepresentation analysis in the vicinity of these uninherited demethylated CpG (uninherited demethylated regions: UDRs) revealed the GATA6 binding motif was overrepresented at 0 h -to- 60 h and 0 h -to- 66 h (Fig. 4D). To further substantiate the overrepresentation of the GATA6 binding motif at the UDRs, we performed ChIPmentation, which can provide evidence for actual physical interactions between genomic regions and GATA6[59]. Consistent with the expression pattern of GATA6, GATA6 binding was not enriched at UDRs during the period 0 h -to- 48 h, indicating the irrelevance of GATA6 during this period (Fig. 4E). In contrast, unlike binding motif overrepresentation, ChIPmentation showed interactions between GATA6 protein and most of the UDRs of all comparisons apart from the 0 h -to- 48 h, consistent with the expression pattern of the GATA6 (Fig. 4E, Fig. S3A). Because ChIPmentation is more direct evidence of TF binding, we assumed that GATA6 binds to the demethylated regions after 48 h. Thus, our results suggest that GATA6 plays a major role in regulating DNA demethylation during DE differentiation.

### The interrelation between DNA demethylation and chromatin status during iPS-DE differentiation

The majority of the demethylated regions were not promoters but other types of regulatory regions such as enhancers and non-annotated regulatory regions (Fig. S4). Therefore, we investigated the chromatin status of the demethylated regions. Active regulatory regions transcribe several classes of transcripts, including mRNA, promoter-upstream transcripts (PROMPTs), and enhancer RNAs (eRNAs), which are typically transcribed within ± 250 bp from the center of the regulatory region[60]. Thus, the transcription level serves as an indicator of chromatin activity. Therefore, to investigate the chromatin activity of the demethylated regions, we measured the average TPM of the UDRs (± 250 bp regions from the uninherited demethylated CpGs) by CAGE. The average TPMs of the UDRs were prone to increase as differentiation proceeds in all comparisons except for the 0 h -to- 48 h, indicating the activation of gene regulatory regions (Fig. 5A).

**Fig. 5.**
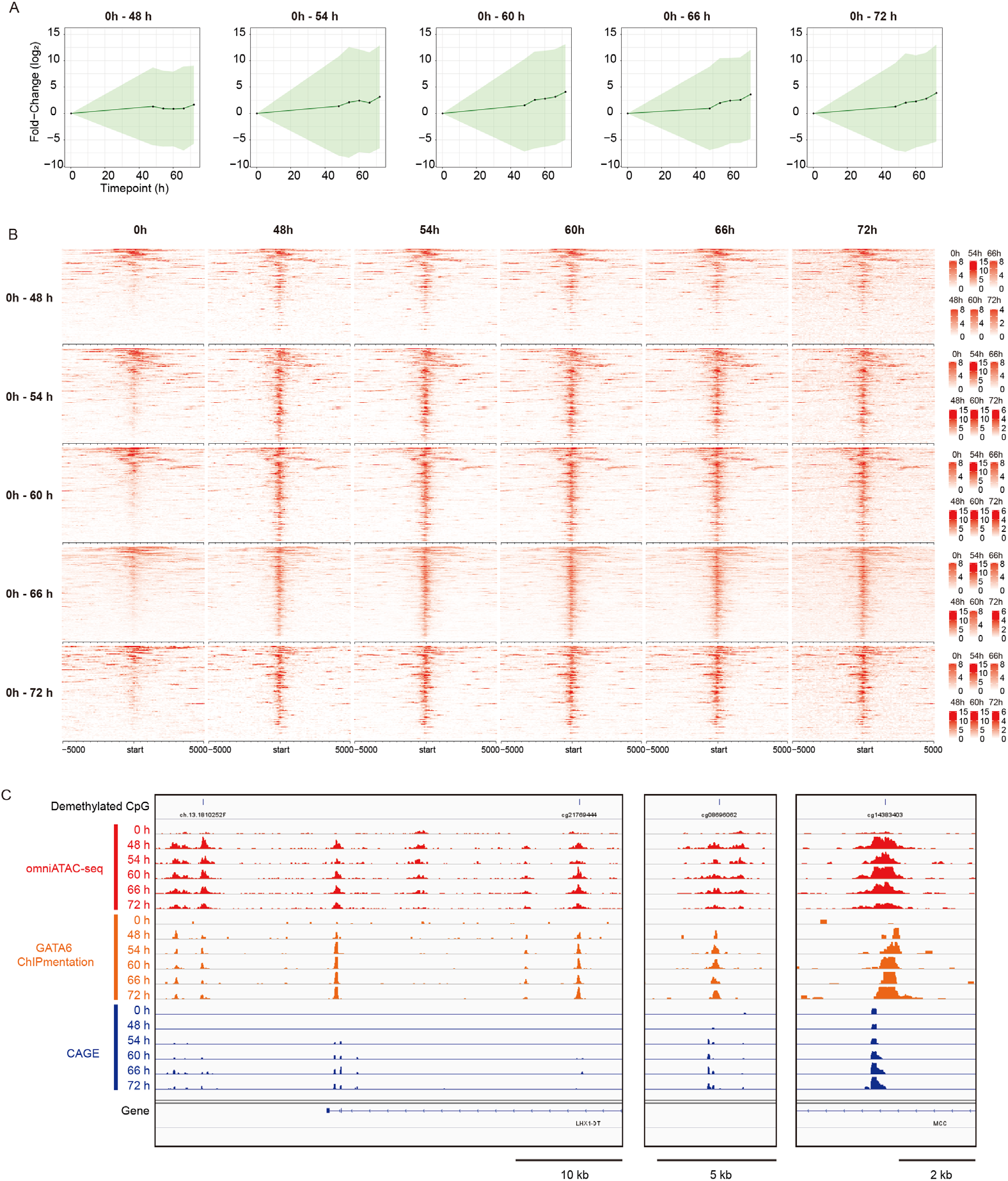
Chromatin status at demethylated regions. **(A)** Change in average TPM of demethylated regions during DE differentiation. X- and Y-axis represents timepoint and relative average TPM (vs. average TPM of 0 h), respectively. The light-green shade is the standard deviation. **(B)** Heatmaps showing Omni-ATAC-seq read coverage at a range of ± 5 kbp from demethylated CpGs. Each time point is horizontally aligned and each of the UDRs are vertically aligned. Red is higher coverage of Omni-ATAC-seq reads. **(C)** A representative screenshot showing DNA demethylated regions, GATA6 ChIPmentation read coverage and OmniATAC-read coverage.

To further analyze the interrelation between GATA6-mediated DNA demethylation and chromatin status, we measured chromatin accessibility by Omni-ATAC-seq[61]. Chromatin accessibility at the UDRs increased between 0 h and 48 h and was maintained over the following timepoints at most of the demethylated regions (Fig. 5B), in agreement with the transcription pattern and GATA6 binding (Fig. 4E, Fig. S2A, Fig. 5B). Notably, the demethylated regions noted during DE differentiation were only marginally accessible in iPS cells (0 h), although GATA6 is not expressed at that time, suggesting that target regions of the GATA6-mediated DNA demethylation are pre-defined by chromatin accessibility (Fig. 5B and 5C).

Taking advantage of our time-course multi-omics dataset, we compared the kinetics of GATA6 expression, GATA6 binding to the genome (ChIPmentation), methylation change (M-value), and chromatin status (ATAC-seq and Transcript) (Fig. 6A). Overall, the kinetics of GATA6 binding, chromatin accessibility, and transcription observed the same trends, regardless of the UDRs. Although GATA6 expression was constantly increasing after 48 h, GATA6 binding plateaued at 54 h, although it was somewhat decreased at 66 h. GATA6 transcription levels increased during 0 h-to-48 h and the expression level was maintained afterward with only slight fluctuations. Of note, chromatin accessibility increased in the period 0 h -to-54 h and then decreased after peaking, in correlation with the methylation change. Thus, the chromatin activation was achieved before the DNA demethylation occurring during DE differentiation.

**Fig. 6.**
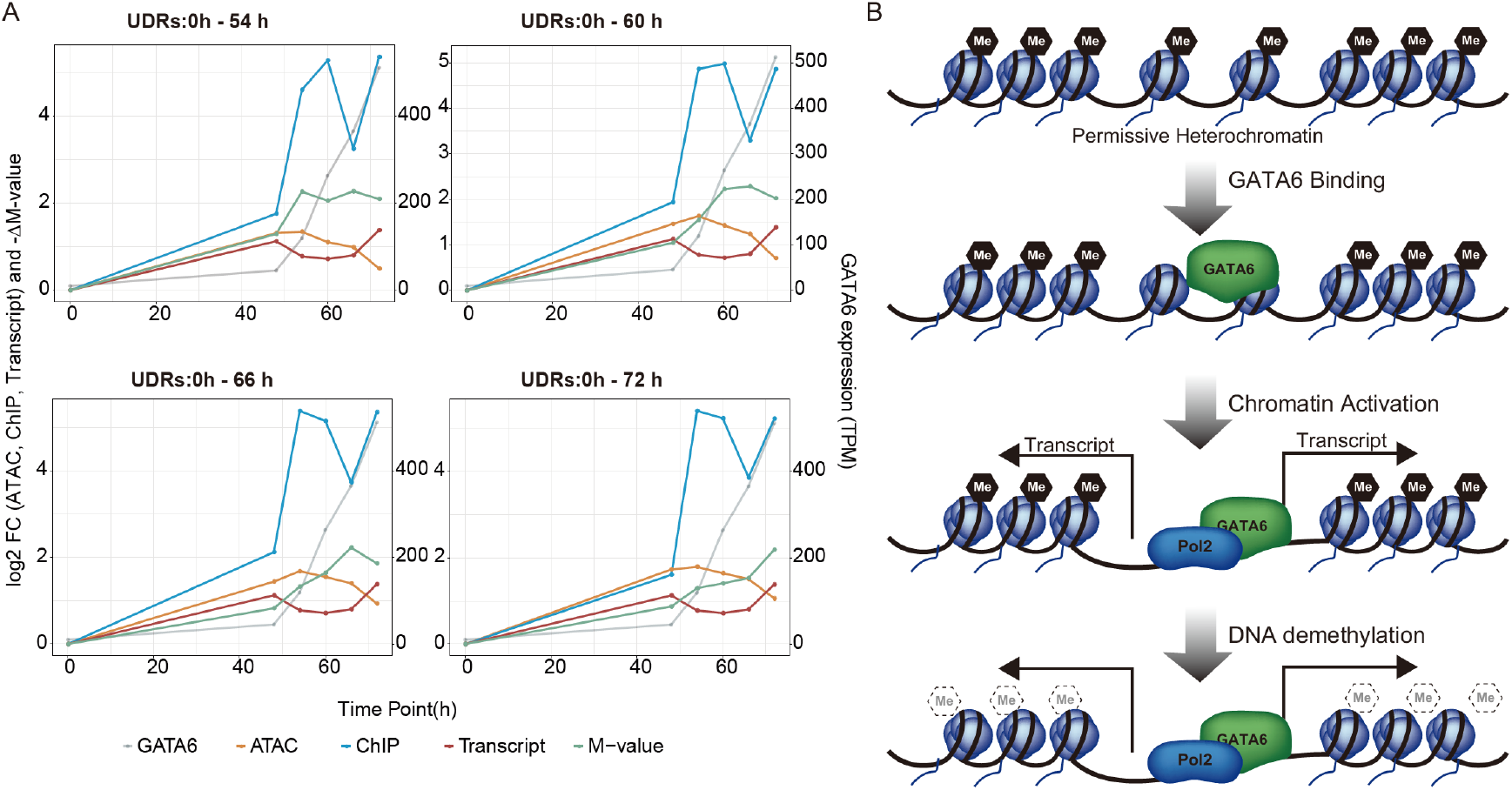
Multi-omics kinetic analysis. **(A)** Line plots showing changes in each demethylated region’s omics data. X-axis is log2 fold-change (FC) for read coverages of ATAC-seq and ChIPmentation for GATA6 (left scale), and -ΔM-value (left scale); TPM for GATA6 expression (right scale). Y-axis represents the time points of the differentiation. **(B)** A schematic illustration showing a model of interrelation between GATA6-mediated DNA demethylation and chromatin status.

## Discussion

In the present study, by applying transcriptome and TFBM overrepresentation analyses for differentially methylated regions, we comprehensively identified putative DNA methylation-regulating TFs active during hepatocyte differentiation. Of these TFs, our results provide multiple strands of evidence that GATA6 is a primary epigenome regulator for the iPSC-to-DE differentiation.

Our data suggest that many TFs participate in modulating DNA methylation dynamics in a stage-specific manner. We previously reported that TF-mediated regulation of DNA methylation predominantly manifests as demethylation[45]. Consistent with that report, we found many TFBMs at demethylated regions during hepatocyte differentiation and the methylation change of these TFBMs tended to be correlated with the expression of corresponding TFs. On the other hand, some TFBMs such as POU5F1 (also known as OCT3/4) were overrepresented mainly at the methylated regions during the iPSC-to-DE differentiation and the expression of the corresponding TFs was inversely correlated with methylation change (Fig. 2B and C; Fig. S2A). In comparison, GATA4 and GATA6 showed binding motif overrepresentation at methylated regions of the hepatoblast differentiation stage when GATA4 and GATA6 expression decreases (Fig. 2B and E). Thus, even the gain of methylation may result from the loss of hypo-methylation maintenance by DNA demethylating-TFs. Interestingly, GATA4 and GATA6 binding motifs are also overrepresented at demethylated regions of the DE differentiation stage when GATA4 and GAT6 expression increases, showing the dual roles of GATA4 and GATA6 (Fig. 2D and E). To summarize, our data suggest that TF-mediated regulation of DNA methylation acts in both the gain and loss of methylation.

Our data also suggest that HNF4A participates in DNA demethylation during the hepatoblast differentiation stage (Fig. 2D). HNF4A is a crucial TF for hepatocyte differentiation and functions, and is reported to be required during liver development for establishment of 5-hydroxymethyl cytosine (5hmC) via interactions with TET3 [3,4,9,62]. An intermediate modification occurring during DNA demethylation, 5hmC has a short half-life and is converted to 5fC and 5caC by TET proteins[36–41]. Then, 5fC and 5caC are rapidly converted to unmodified cytosines by base-excision repair. Because the methylation array analyses used in the present study do not distinguish between methylated cytosine and 5hmC, our results suggest that HNF4A-induced 5hmC is immediately converted to unmodified cytosine.

Out of the putative DNA-demethylating TFs that we identified, our data demonstrated that GATA6 plays a pivotal role in DNA demethylation during DE differentiation. GATA6 mRNA expression started increasing at 48 h and was dramatically upregulated during the DE differentiation stage (Fig. 3A and Fig. S2A). In parallel with the expression, binding of GATA6 proteins to the vast majority of demethylated regions was detected and this was maintained through differentiation (Fig. 4E, Fig. 6A), suggesting GATA6 promotes DNA demethylation at its binding sites. In support of this molecular function of GATA6, ectopic expression of GATA6 in HEK 293T cells proved GATA6-mediated binding site-directed DNA demethylation (Fig. 3C). Thus, these results demonstrate that GATA6 is a crucial regulator of DNA demethylation for early hepatic development, and acts in a binding site-directed manner.

In the analysis of the iPS cells -to- DE time-course, GATA6 binding motif overrepresentation was not consistent with the results from ChIPmentation for GATA6. Although ChIPmentation showed GATA6 protein binding at the UDRs of all comparisons except 0h -to- 48 h, binding motif overrepresentation was only detected at the UDRs of the 0 h -to- 60 h and 0 h-to- 66 h comparisons. For the GATA6 binding motif overrepresentation analysis, we used the GATA6 PWM of the IMAGE motif database, which includes the canonical GATA binding motif GATW (W = A or T). However, GATA-binding proteins can bind various motifs that differ from the canonical GATA-binding motif with comparable affinities [63]. Therefore, TFBM overrepresentation analysis using a known motif database may underestimate the TF binding. Another possibility is that ChIPmentation includes indirect binding of GATA6 via their co-factors. For instance, Friend Of GATA (FOG) proteins, which are co-factors of GATA proteins, have been reported to play essential roles in mediating DNA loop formation[64]. Thus, TFBM overrepresentation analysis with a known motif may not be completely reflecting actual TF binding. Nevertheless, TFBM overrepresentation has a value in predicting the TF binding because it is only based on *in-silico* analysis without experimental fluctuation. It is also noteworthy that ChIPmentation and ChIP-seq depend highly on the quality of the antibody, which often leads to experimental unreliability.

We also found a relationship between GATA6-mediated DNA demethylation and chromatin activation. Notably, chromatin accessibilities of GATA6 binding regions are already slightly accessible in iPS cells (Fig. 4E), although GATA6 is not expressed at that stage. GATA6 appears to be a pioneer factor that directly binds to permissive heterochromatin and primes the opening of chromatin [65–69]. Consistent with this, our result indicates that the targets of GATA6-mediated DNA demethylation are preliminarily marked by marginal chromatin accessibility. Thus, the chromatin accessibility assay preliminarily indicates the target regions for TF-mediated regulation of methylation.

Further multi-omic kinetics analysis suggested the temporal relationships that exist between GATA6-mediated DNA demethylation and chromatin activity. Interestingly, chromatin accessibility increased from 0 h -to- 54 h and then declined afterward, although DNA methylation decreased. This is inconsistent with the notion that DNA methylation is correlated with closed chromatin. The methyl-group of methylated cytosine lies in the major groove of the DNA double helix, which hinders the interaction of TFs with DNA. On the other hand, DNA demethylation increases the affinity of the TFs for their binding site. Therefore, the decrease in chromatin accessibility may be due to occupation of the opened chromatin by TFs. In fact, the chromatin accessibility assay reflects not only the presence of open chromatin or nucleosome density but also TF binding[70]. Thus, our results indicate that the chromatin accessibility assay may not correctly reflect the chromatin activity.

Although the underlying molecular mechanisms have not been investigated in this study, our analysis proposes a sequential reaction takes place, coordinated with the expression pattern of TFs. DNA-demethylating TFs firstly bind to the permissive heterochromatin sites where the TFBM are located. They then open and activate the chromatin at the binding sites, and finally complete DNA demethylation (Fig. 6B). This sequential reaction may be due to differences in reaction times between chromatin remodeling and DNA demethylation, because the level of DNA methylation progressively decreases from the beginning of the differentiation process. While chromatin remodeling is an enzymatic reaction, DNA demethylation is achieved by several mechanisms, including passive DNA demethylation, which depends on cell division. Cell division is a complex process composed of multiple steps, and taking more time than a single enzymatic reaction. Therefore, even if timing for the initiation step is the same, the total reaction time to completion may differ between chromatin remodeling and DNA demethylation.

In addition to GATA6 having an essential role in physiological endoderm cell development, GATA6 haploinsufficiency causes several diseases such as neonatal diabetes mellitus, cardiomyopathy, and pancreatic agenesis[71–73]. In the present study, we found a novel function of GATA6, regulating binding site-directed DNA demethylation. Hence, epigenetic abnormalities may also be associated with the pathology of these diseases already linked to GATA6. Hence, epigenetic analyses of these diseases deserves to be a priority and may provide novel insights into underlining molecular mechanisms.

## Conclusions

We identified multiple putative DNA methylation-regulating TFs acting at distinct stages throughout hepatocyte differentiation, which are likely involved in DNA demethylation and maintenance of hypo-methylation. Our data suggest that multiple TFs cooperatively modulate the DNA methylation landscape during cellular differentiation. A finer scale analysis of the time-course throughout DE differentiation showed the crucial role of GATA6-mediated DNA methylation regulation, which is gradually completed upon the rapid activation of chromatin.

## Methods

### Cell culture and in vitro differentiation

The 201B7 human iPS cell line was acquired from the RIKEN BioResource Center (BRC) and was cultured in a Cellartis® DEF-CS™ Culture System (Takara Bio Inc., Shiga, Japan). For *in vitro* hepatocyte differentiation and DE differentiation, we used the Cellartis® Hepatocyte Differentiation Kit (Takara Bio Inc.) and the Cellartis^®^ DE Differentiation Kit (Takara Bio Inc.), respectively, according to the manufacturers’ instructions.

### Methylation array analysis

Genomic DNA was isolated using a NucleoSpin® Tissue Kit (Macherey-Nagel, Düren, Germany). The methylation array used an Infinium Human methylationEPIC BeadChip (Illumina, San Diego, CA), according to the manufacturer’s instructions. Data was processed as previously described.

### Cap Analysis Gene Expression

Total RNA was extracted using NucleoSpin® RNA (Macherey-Nagel). CAGE libraries were prepared as previously described. Briefly, 3 μg of total RNA from each sample were used in reverse transcription reactions with random primers. The 5’ end cap structure was biotinylated and captured with streptavidin-coated magnetic beads (Thermo Fisher). After ligation of 5’ and 3’ adaptors, second-strand cDNA was synthesized, followed by digestion with exonuclease I (New England BioLabs). The purified CAGE libraries were sequenced using single-end reads of 50 bp on the Illumina HiSeq 2500 (Illumina, USA). The extracted CAGE tags were then mapped to the human hg19 genome by STAR. The tags per million (TPM) were calculated for each FANTOM5 TSS peak and regions extended ± 250 bp from each differentially methylated CpG. Gene expression levels of each gene were computed as the sum of multiple TSS peaks associated with a single gene.

### Omni-ATAC-seq

Omni-ATAC-seq libraries were prepared as previously described[61]. Briefly, 5 × 10^4^ cells were stored at −80 °C in STEM CELLBANKER® (Takara Bio Inc.) until use. The cells were washed with PBS and nuclei were extracted. The extracted nuclei were resuspended in 50 μl of transposition mix (100 nM TED1 (Illumina), 0.01% digitonin, and 0.1% Tween-20, in TD buffer (Illumina)) and incubated at 37 °C for 30 min with 1,000 RPM mixing. DNA was extracted from the reaction mixture with Zymo DNA Clean and Concentrator (Zymo Research, CA, USA). DNA library was prepared using NEBNext® Ultra™ DNA Library Prep Kit for Illumina® (New England BioLabs) with 5cycles of pre-amplification and 3 to 7 cycles of PCR amplification. Amplified DNA library was purified with Zymo DNA Clean and Concentrator (Zymo Research), followed by two size-selection steps with SPRIselect (1:0.6 and 1:0.2 sample vol. to beads vol.; Beckman Coulter, CA, USA). The libraries’ size distribution was determined by Bioanalyzer (Agilent Technologies, CA, USA), and the concentration of the libraries was quantified by GenNext NGS Library Quantification Kit (Toyobo Co., Ltd., Osaka, Japan). The Omni-ATAC-seq libraries were sequenced using 150 bp paired-end reads on the HiSeq X (Illumina). The obtained sequence reads were mapped to the human hg19 genome by bowtie2.

### Lentivirus preparation and transduction

GATA6 and ORF were sub-cloned into the CSII-EF-RfA-IRES2-puro vectors using the Gateway LR reaction (Thermo Fisher Scientific Inc.). GATA6 lentivirus vectors were produced by using the LV-MAX Lentiviral Production System (Thermo Fisher Scientific Inc.) according to the manufacturer’s instructions. The resulting lentivirus vectors were transduced to 293T cells, as described previously.

### Quantitative reverse transcription PCR (qRT-PCR)

qRT-PCR was performed as previously described[45] with primers shown in table S1.

### ChIPmentation

ChIPmentation was performed using a ChIPmentation for Transcription Factor kit (Diagenode) according to the manufacturer’s instructions. Briefly, the cells were collected and fixed with 1% formaldehyde for 8 minutes at RT. The fixed cells were lysed, and chromatin was sheared by sonication using a Picoruptor® (Diagenode) for 10 cycles. The sheared chromatin derived from one million cells was subjected to magnetic immunoprecipitation and tagmentation using an SX-8G IP-STAR® Compact Automated System (Diagenode) with the anti-GATA6 antibody (D61E4, Cell Signaling Technology, Inc.). The immunoprecipitated samples were stripped from the magnetic beads and subjected to end repair and reverse cross-linking. The Illumina sequence compatible sequencing libraries were amplified by nine cycles of PCR. The sequencing libraries were cleaned up using AMPure XP beads (1:1.8 sample vol. to beads vol.; Beckman Coulter). The size distribution of the libraries were determined by Bioanalyzer (Agilent Technologies). The concentration of the libraries was quantified by GenNext NGS Library Quantification Kit (Toyobo Co., Ltd). The ChIPmentation libraries were sequenced using 150 bp paired-end reads on the HiSeq X (Illumina). The sequence reads that were obtained were mapped to the human hg19 genome by bowtie2.

## Computational Methods

### Functional analysis of differentially methylated regions

GO analysis of differentially methylated regions was performed using *GREAT*[53]. Enriched GO lists were summarized based on Semantic Similarity by the GOsemSim R package.

### Screening of DNA methylation-regulating transcription factors

TFBM overrepresentation analysis was performed as previously described with an additional modification. Briefly, sequences located ± 5 kbp from the methylated or demethylated probe positions and the same number of randomly selected probes were extracted from version hg19 of the human genome sequence. TFBM identification was performed using the *matchPWM* command of the Biostrings package of Bioconductor with the PWM database of Integrated analysis of Motif Activity and Gene Expression changes of transcription factors (IMAGE). Out of the overrepresented motifs, the corresponding genes whose CAGE tag per million ≥ 50 at the time points where the TF binding motif was overrepresented were selected as DNA methylation-regulating transcription factors.

### Correlation matrix

The correlation coefficient of all combinations of two clusters was computed using the M-values. The correlation coefficients were visualized as the correlation matrix heatmap. The clusters were ordered based on hierarchical clustering, which was calculated using the *hclust* and *dist* functions of the R stats package with the default settings.

### Functional analysis of differentially methylated regions

Differentially methylated CpGs that were identified as ΔM > 2 and ± 100 bp extended regions from the differentially methylated CpGs were used as differentially methylated regions. The differentially methylated regions were subjected to GREAT analysis using the *submitGreatJob* function implemented in the rGREAT R package with background data, which is with the regions extended ± 100 bp for all methylation array probes. Log_10_ FDR and ratio between the numbers of hit regions and all differentially methylated regions of the Top10 overrepresented GOs (Biological Process) were visualized.

### Annotation of differentially methylated regions

Gene promoters were defined as 1 kbp upstream and 200 bp downstream regions of genes in gencode human release version 19. The enhancers used in this study were FANTOM 5 human phase 1 and 2 permissive enhancers. Non-promoter and non-enhancer regions were defined as unannotated regions. The complete overlap between uninherited demethylated CpGs and each regulatory region was counted.

### Coverage analysis of GATA6 ChIPmentation and Omni-ATAC-seq

Bigwig Coverage files of CAGE and Omni-ATAC-seq were computed using *bam2wig.py*. The read coverage was visualized in the range between ± 5 kbp from the demethylated CpGs using the *EnrichedHeatmap* function implemented in the EnrichedHeatmap R package.

## List of abbreviations

TFs: transcription factors
TFBM: transcription factor binding motif
PBS: Phosphate-buffered saline
DE: definitive endoderm
UDRs: uninherited demethylated regions

## Declarations

### Ethics approval and consent to participate

Not applicable.

### Consent for publication

Not applicable.

### Authors’ contributions

TS participated in the study’s design, devised the methodology, performed the statistical analyses, carried out the molecular biology studies, acquired the funding, and drafted the manuscript. SM, EF, MK, YM, YT, JL, HN, SA, and YS carried out the molecular biology experiments. HS helped to draft the manuscript, acquired the funding, and supervised the study. All authors read and approved the final manuscript.

## Acknowledgments

We thank Chung-Chau Hon for useful advice in data analysis. We thank Jing-ru Li for experiment supports. We thank Horoyuki Miyoshi and RIKEN BRC for providing lentivirus plasmids. We are grateful to RIKEN IMS, Laboratory for Comprehensive Genomic Analysis, for the Hiseq 2500 sequencing.

