## Supplementary Material Legends for "Master Transcription Factors Regulate the DNA Methylation Landscape During Hepatocyte Differentiation"

**Fig. S1 Expression of differentiation marker genes.** Line plot showing average TPM of each promoter of pluripotent, DE, and hepatic differentiation marker genes. The error bar is the standard deviation. X- and Y-axis are days of differentiation and average TPM, respectively. The experiment was done in triplicate. (B, C) Top 5 enriched summarized GOs of methylated (B) and demethylated (C) regions. The dot size represents the ratio between the number of probes that hit the GOs and all methylated or demethylated probes. The color represents total scores computed by *reduceSimMatrix* function of the rrvgo R package.

**Fig. S2 Expression profile of putative DNA methylation-regulating TFs**
(C, E) mRNA expression profile of the genes corresponding to the overrepresented TF binding motifs at methylated (C) and demethylated (E) regions. X- and Y-axes show time points of differentiation (hours from differentiation initiation) and tag-per-million (TPM) of CAGE, respectively. The color of each line represents the maximum TPM.

**Fig. S3 Transcriptome and methylome analysis of iPS cell -to- DE differentiation.**
(A) Line plot showing average TPM of each promoter of the pluripotent, primitive streak, and DE differentiation marker genes. The error bar is the standard deviation. X- and Y-axis are days of differentiation and average TPM, respectively. The experiment was done in triplicate. (B, C) Distribution of enrichment score for the GATA6 binding motif within ±5,000 bp of demethylated CpG probes at each interval of adjacent timepoint (B) and 0 h vs. 72 h (C). X- and Y-axes show distance from probe CpG position and enrichment score, respectively. Horizontal and vertical lines are enrichment score = 0 and demethylated CpG position, respectively. The colors of each plot represent colors of timepoints shown in Fig. 4A. (D) A heatmap showing the M-value at each timepoint of the probes demethylated in 72 h compared with 0 h. The color represents the z-scored M-value.

**Fig. S4 The proportion of UDRs by regulatory region annotation.** X- and Y-axis represent comparison and proportion, respectively.

**Table S1 Primer list**
