## Supplementary figures and images for "Master Transcription Factors Regulate the DNA Methylation Landscape During Hepatocyte Differentiation"

### Supplementary Figure 1

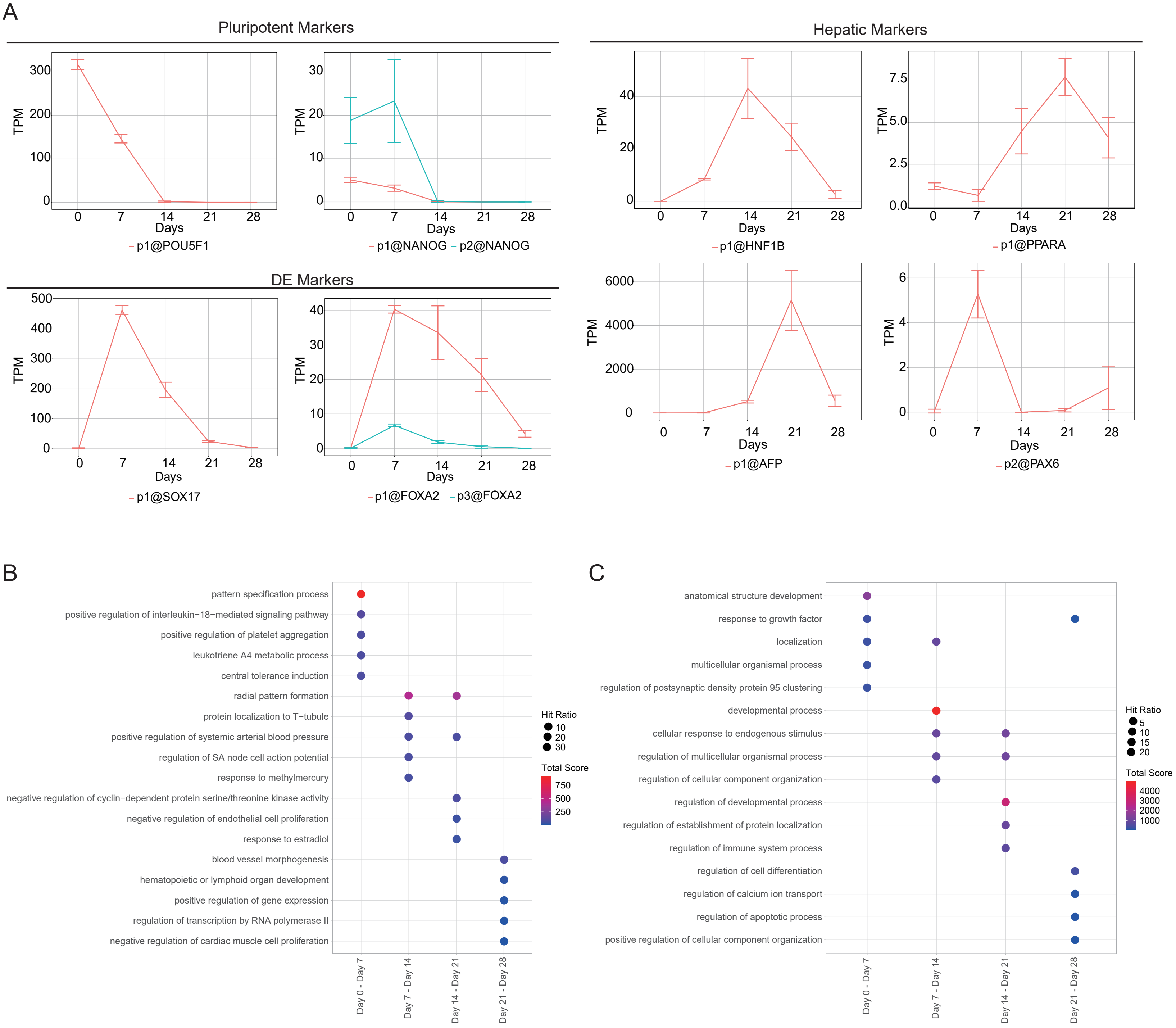

### Supplementary Figure 2

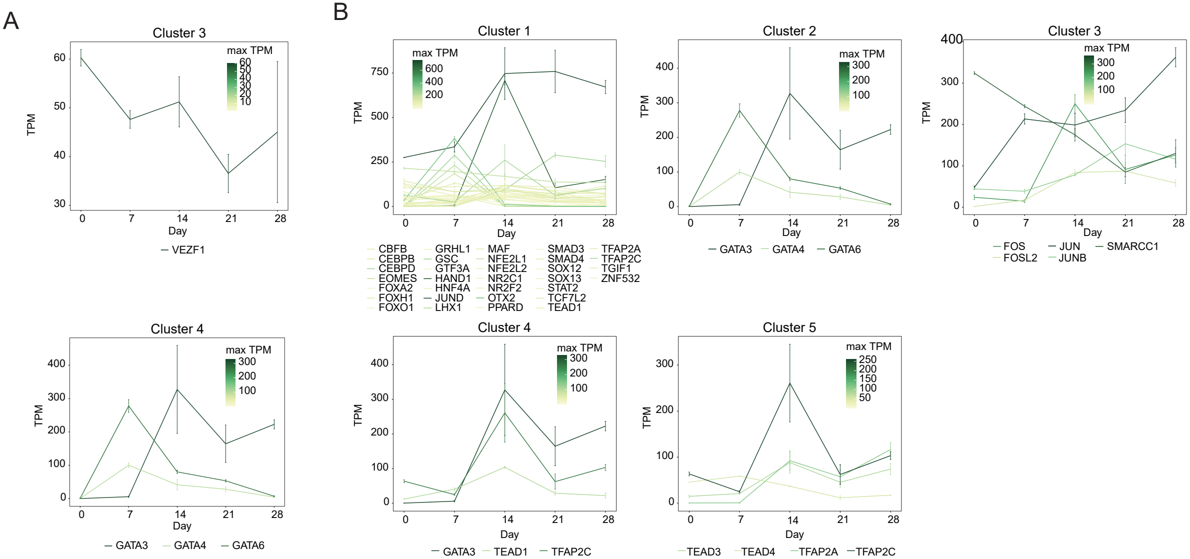

### Supplementary Figure 3

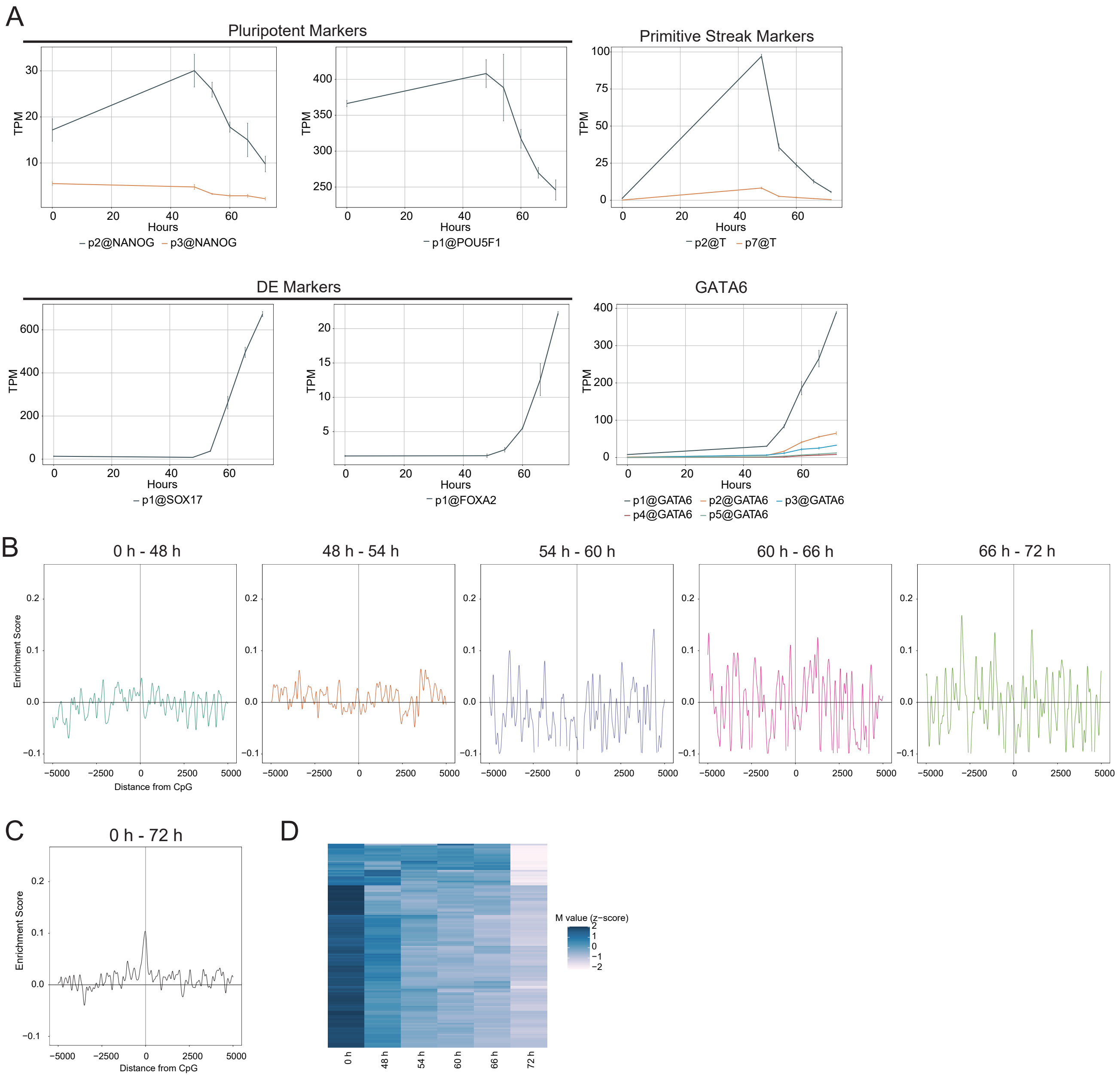

### Supplementary Figure 4

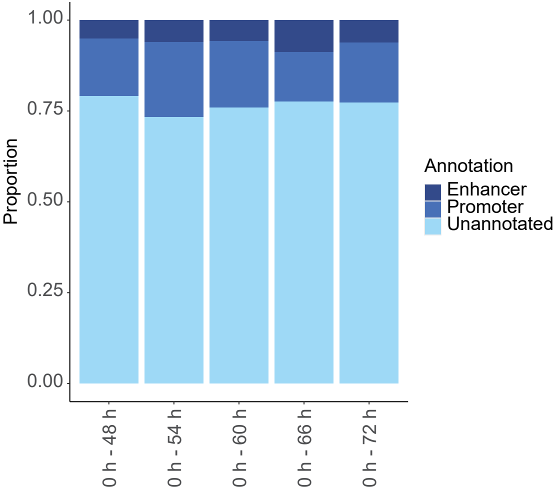
